# Engineered Transdermal Peptide-Recombinant Type III Collagen Hydrogel with Biological Efficacy in Cell Proliferation and Wound Healing

**DOI:** 10.1101/2025.11.22.689886

**Authors:** Wansen Tan, Yue Liu, Jingjun Hong

## Abstract

Collagen, as a major structural protein in the extracellular matrix, plays a crucial role in tissue regeneration; however, traditional collagen sources often suffer from immunogenicity, poor stability, and batch-to-batch variability. In this study, a novel transdermal peptide-recombinant type III collagen hydrogel was developed, which has excellent biocompatibility, thermal stability, and skin repair effects. Transdermal peptide-recombinant type III collagen was expressed and purified by genetic engineering methods, and its potential for biomedical applications was further evaluated. The experimental results showed that the prepared collagen had a high purity (95%) and retained the unique secondary structure of collagen, showing good structural stability at different pH values. Through cell proliferation experiments and mouse wound healing experiments, we verified the superior effect of the collagen hydrogel in promoting wound healing and significantly accelerating wound healing at concentrations as low as 0.2 mg/mL. This achievement provides strong experimental support for its application in clinical skin wound repair.

## 1. Introduction

With the continuous advancement of biomaterial research, especially in the fields of regenerative medicine and tissue engineering, collagen, as a natural biomaterial, is widely used in trauma treatment, skin repair, and various regenerative medicine products due to its good biocompatibility and ability to promote cell growth and tissue repair[1; 2]. There are many types of collagen, among which type III collagen is an important component of the skin, blood vessels, and other connective tissues, and its main function is to provide structural support and participate in the wound healing process[3]. Therefore, type III collagen has important application prospects in tissue engineering and skin wound repair.

However, traditional sources of type III collagen usually come from animal tissues, especially cattle, pigs and other animals. However, this source presents several issues.[4; 5]. First, collagen of animal origin may carry potential pathogenic bacteria, viruses or other pathogenic factors, which may cause immune reactions or even allergic reactions during use. Secondly, obtaining sufficient amounts of high-quality collagen faces challenges due to limitations in animal resources, including production costs and ethical concerns. To overcome these limitations, in recent years, scientists have adopted genetic engineering techniques to express high-quality collagen in microbial or mammalian cells through recombinant DNA technology[6–8], thereby achieving its mass production and reducing costs.

Significant advances have been made in genetically engineered expression of type III collagen, and many studies have focused on developing recombinant collagen with excellent biological properties. However, existing recombinant collagen products often face problems such as poor stability, structural instability, and poor use effect, especially in the application of hydrogels, which are particularly prominent[9–12]. Hydrogel, as a highly hydrated and softened biomaterial, has been widely used in wound dressings and skin repair agents. However, traditional hydrogels are susceptible to environmental conditions and lose their structural stability during long-term application, leading to collagen degradation and thus affecting the repair effect[13–16].

Therefore, this study aims to develop a novel transdermal peptide-recombinant type III collagen[17] hydrogel that expresses recombinant type III collagen through genetic engineering technology and combines it with a hydrogel carrier to improve its biological activity and application effect in skin repair. Through detailed analysis of the physicochemical properties, molecular structure, and stability of the collagen under different pH conditions, we found that the protein has good thermal and structural stability, and can maintain its secondary structure and adapt to the physiological environment of the skin. In addition, we have verified the superior biological activity of the hydrogel material through a series of in vitro and in vivo experiments, including cell proliferation promotion and accelerated wound healing.

In particular, we found that the hydrogel can significantly promote the healing of wounds in mice at low concentrations (0.2 mg/mL), and has good gender adaptation, showing a faster repair effect than traditional collagen dressings. Compared to traditional animal-derived collagen dressings, recombinant collagen hydrogels not only avoid potential immune responses and the risk of pathogen transmission, but also improve the sustainability and cost-effectiveness of production. Due to its excellent biocompatibility and stability, it has a wide range of clinical application potential, especially in the fields of wound repair, skin care, and regenerative medicine.

In general, the transdermal peptide-recombinant type III collagen hydrogel developed in this study provides a new idea for the application of collagen-like materials, which not only overcomes the shortcomings of traditional collagen sources, but also greatly improves the application value of hydrogels in clinical treatment. It is hoped that this research result can promote the industrialization of collagen-based biomaterials and provide a safer and more effective option for future wound repair treatment.

## 2. Results and discussion

### 2.1 Physicochemical Properties of Transdermal Peptide–Recombinant Type III Collagen

An engineered Pichia pastoris strain was constructed to express a transdermal peptide-recombinant type III collagen fusion protein. The recombinant protein was purified from the fermentation broth using ion-exchange chromatography and subsequently characterized for molecular weight and purity. Following purification, the molecular weight and purity of the transdermal peptide-recombinant type III collagen were analyzed (Figure 1). Size-exclusion chromatography (Superdex^TM^ 200 Increase 10/300 GL) (Figure 1A) revealed that the protein forms a stable trimer. This was consistent with the molecular mass of 21,477 Da determined by mass spectrometry (Figure 1B). SDS-PAGE analysis showed an apparent molecular weight of approximately 25 kDa, appearing as a single band (Figure 1C). High-performance liquid chromatography (HPLC) analysis further demonstrated that the purified protein exhibited a purity of 95% (Figure 1D). The purified transdermal peptide-recombinant type III collagen exhibits high purity and conforms to the expected molecular weight.

**Figure 1.**
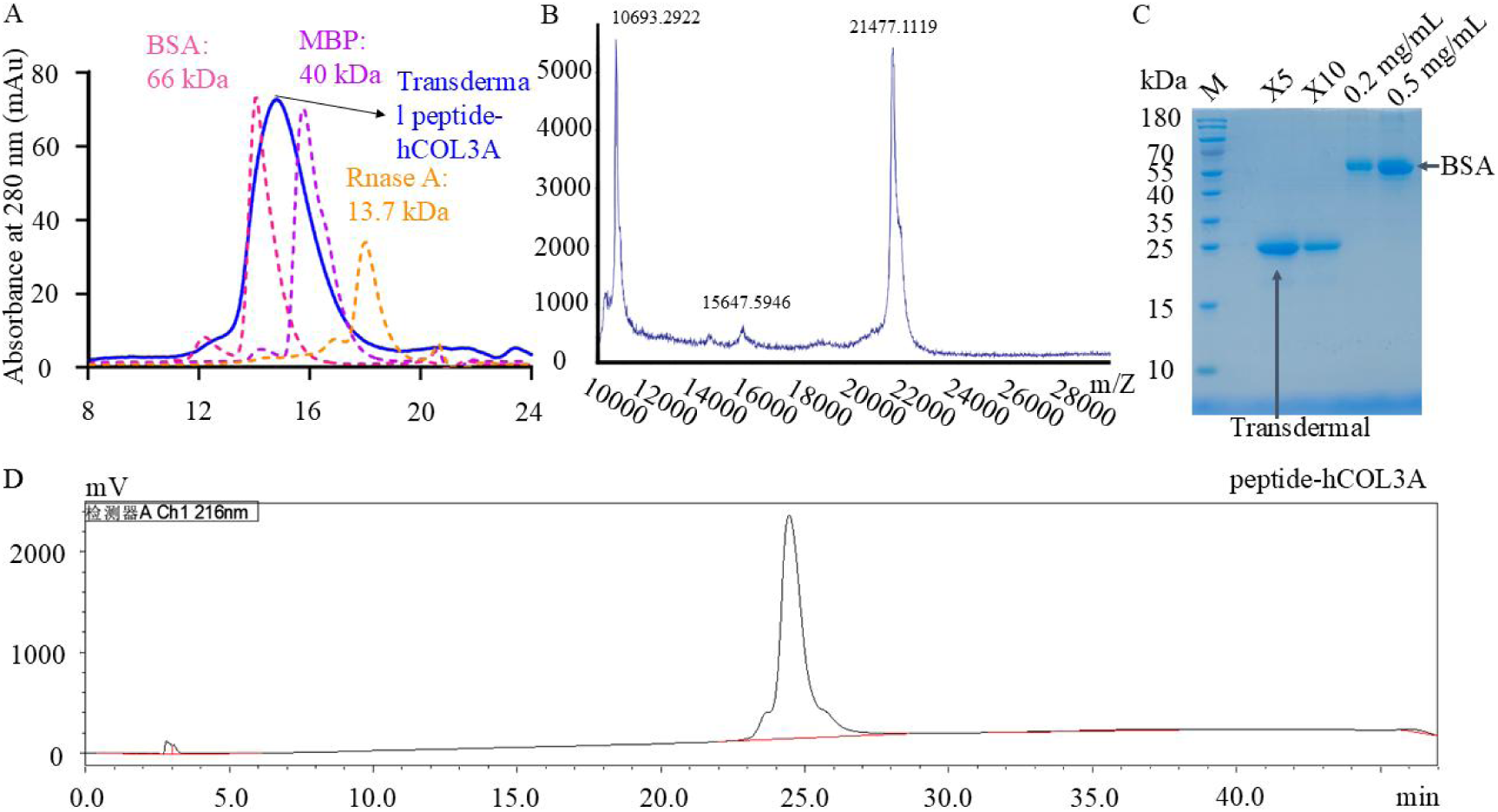
Purity and molecular weight analysis of transdermal peptide–recombinant type III collagen. (A) Size-exclusion chromatography indicates that transdermal peptide recombinant type III collagen exhibits a molecular weight between MBP and BSA, confirming its trimer. (B) Mass spectrometry (MALDI-TOF) measurement confirming the molecular mass at 21,477 Da. (C) SDS-PAGE analysis of the recombinant collagen revealed a single band with an apparent molecular weight of approximately 25 kDa. (D) High-performance liquid chromatography (HPLC) profile indicating a purity of 95% for the recombinant collagen.

The secondary structure of the recombinant protein was further analyzed by circular dichroism (CD) spectroscopy. In both pH 6.2 (Figure 2A) and pH 7.2 (Figure 2B) buffers, the protein exhibited a pronounced negative peak near 197 nm, characteristic of the left-handed poly(L-proline) type II helix, a hallmark of collagenous structures. This spectral signature remained unchanged between pH 6.2 and 7.2, demonstrating the protein’s conformational stability across physiologically relevant conditions. The structural robustness observed under these mildly acidic to neutral pH levels confirms that the recombinant type III collagen can maintain its native fold in the presence of common hydrogel excipients. These findings collectively affirm the structural stability of the transdermal peptide recombinant type III collagen, supporting its suitability for hydrogel formulation and broader biomedical applications.

**Figure 2.**
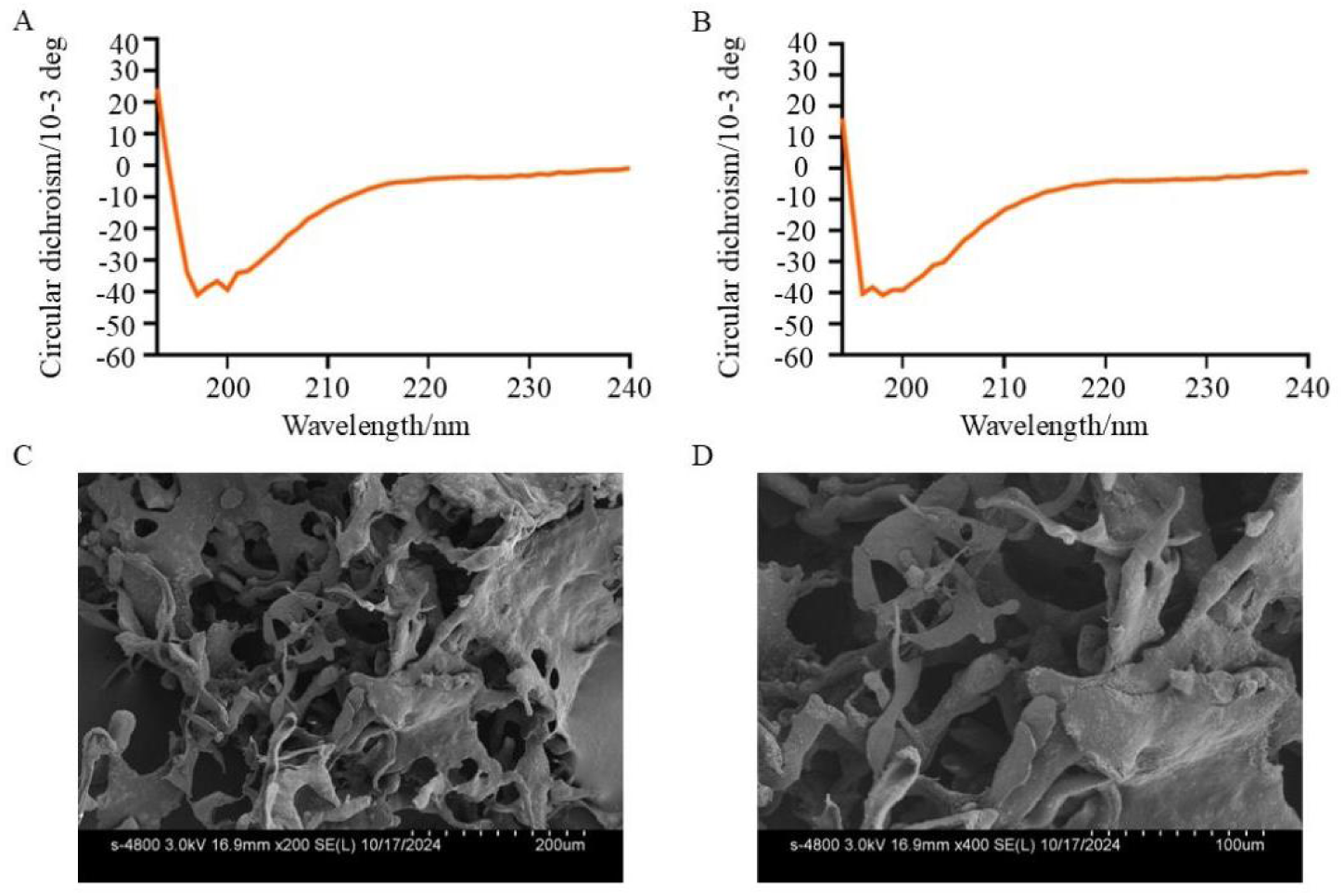
Circular dichroism (CD) spectra of transdermal peptide–recombinant type III collagen. (A) CD spectrum at pH 6.2, displaying a characteristic negative peak near 197 nm. (B) CD spectrum at pH 7.2, demonstrating structural consistency and conformational stability under physiologically relevant pH conditions. (C) Scanning electron microscopy (SEM) image of lyophilized recombinant collagen at 200× magnification, revealing a lamellar, network-like architecture. (D) SEM image at 400× magnification, highlighting the rough surface topography and abundant porous structure of the protein.

The microstructural morphology of lyophilized recombinant collagen was examined by scanning electron microscopy (SEM) at 200× (Figure 2C) and 400× magnification (Figure 2D). The protein displayed a lamellar, interconnected network structure with abundant porous regions and a rough surface topography, features that confer potential for mechanical support in biomedical applications.

### 2.2 Biological activity of transdermal peptide recombinant type III Collagen

To confirm that the safety of transdermal peptide recombinant type III collagen-the main raw material of the hydrogel — met the required standards and its biological activity was better than that of recombinant type III collagen, which was commonly used at present, we conducted in vitro metamorphic skin reaction assays, transdermal detection and cell proliferation experiments.

In vitro allergy skin reaction assay. The erythema and edema of guinea pig skin after administration were observed by stimulation of the skin surface of 300 g∼500 g of newly adult healthy guinea pigs (Figure 3), combined with Table 1, there was no erythema edema on the skin surface of guinea pigs after the first dose and the second dose stimulation 2 weeks later, which met the safety requirements. Further, we used 6∼8 weeks of Kunming rats to conduct transdermal experiments on female and male rats, comparing the cumulative transdermal amount between transdermal peptide recombinant type III collagen and non-transdermal peptide recombinant type III collagen. Measurement of cumulative transdermal amount after 8 hours showed that the transdermal peptide-modified version of recombinant type III collagen outperformed the standard one by 43.6% in male and 42.8% in female mouse skin. We found that both female and male mice showed superior transdermal absorption of the peptide–type III collagen (Figure 4).

**Figure 3.**
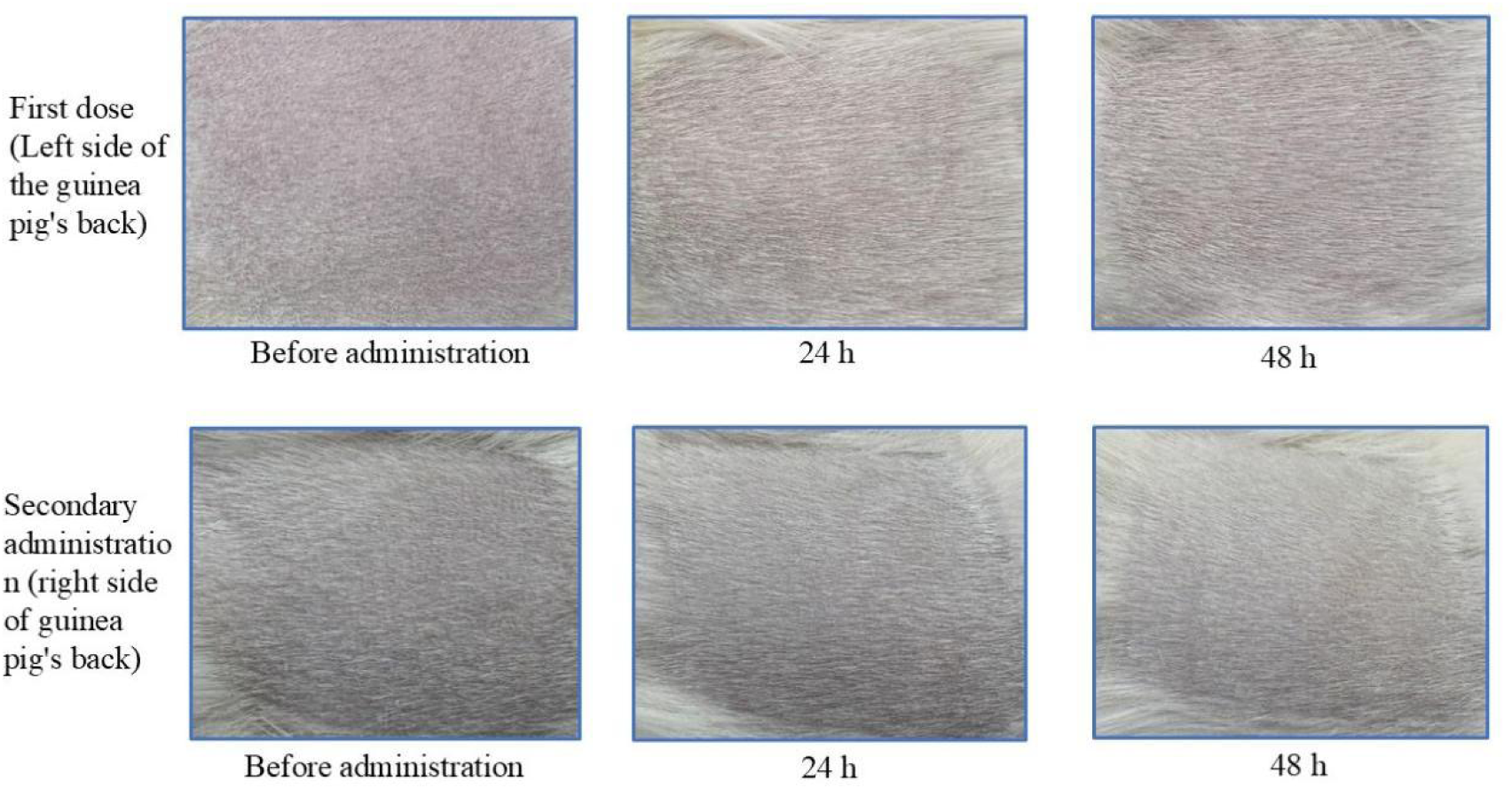
In vitro skin reaction assay in guinea pigs after administration of transdermal peptide recombinant type III collagen. No erythema or edema was observed after both the first and second dose, indicating safety for topical use.

**Figure 4.**
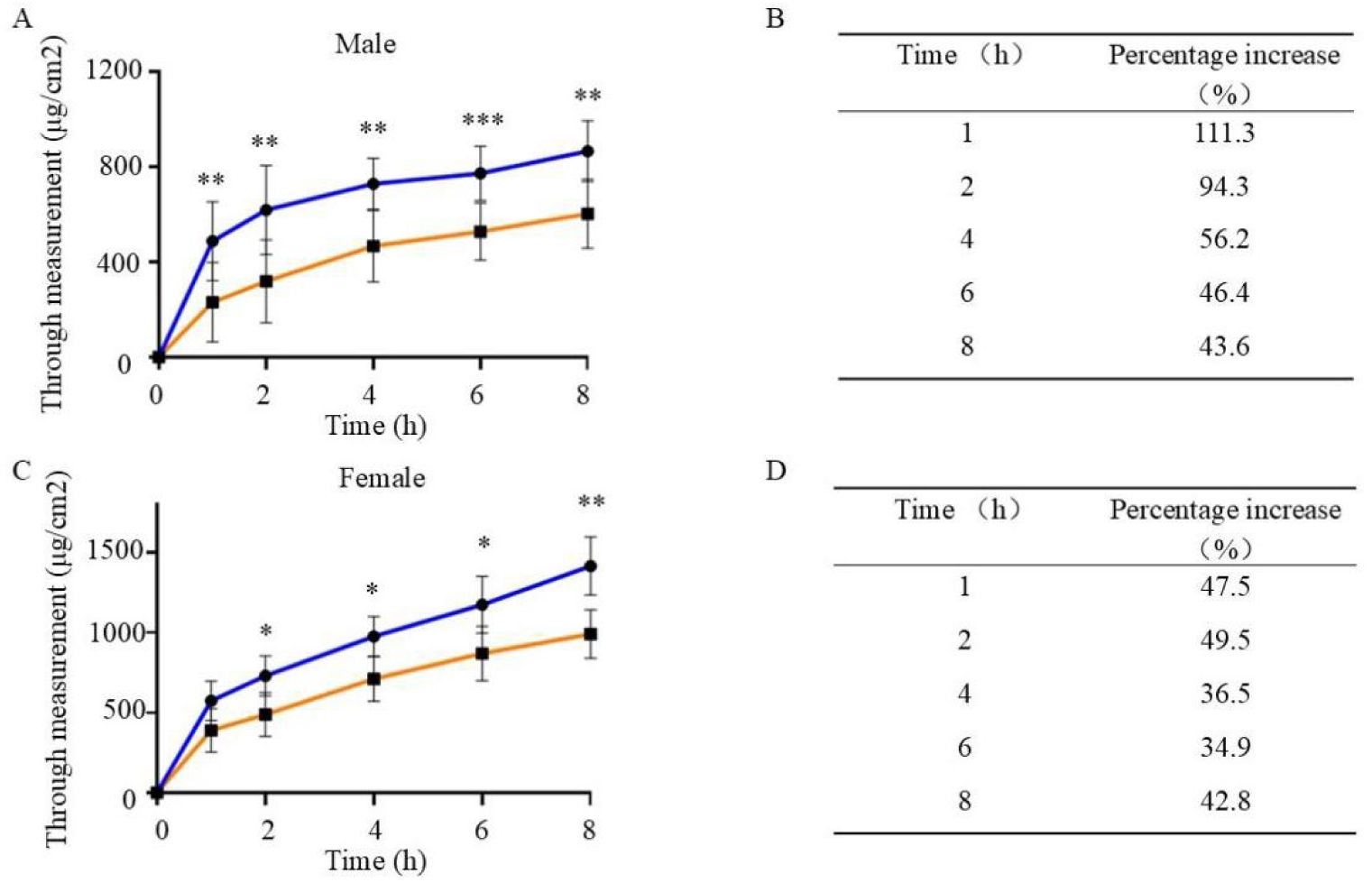
Permeation experiment of recombinant type III collagen on excised mouse skin between female and male (Transdermal peptide recombinant type III collagen - blue, non-transdermal peptide recombinant type III collagen – orange; The data underwent T-test significance analysis, and its specific star-level correspondence is ⁎: 0.01<P<0.05, ⁎⁎: P<0.01). (A) Analysis of the permeability and significance of collagen on the skin of male mice in vitro at 0∼8h. (B) Increased rate of transdermal peptides on the ex vivo skin of male mice. (C) Analysis of the transmittance and significance of collagen on the skin of female mice in vitro at 0∼8h. (D) The rate of increase brought by transdermal peptides on the ex vivo skin of female mice.

**Table 1:** Cosmetic Safety Technical Specifications - In Vitro Allergic Skin Reaction Scoring Table.

| Scoring Table |  |
| --- | --- |
| Skin reactions | Integral |
| No erythema | 0 |
| Slightly erythema | 1 |
| Obvious erythema | 2 |
| Moderate-severe erythema | 3 |
| Severe erythema | 4 |
| No edema | 0 |
| Mild edema | 1 |
| Moderate edema | 2 |
| Severe edema | 3 |

In the cell proliferation experiment, HaCaT cells were cultured by adding recombinant type III collagen raw materials from different sources, and the cell viability of HaCaT cells after 24 hours of culture was determined by CCK-8 reagent, and it was found that the recombinant type III collagen concentrations were 0.025, 0.05, and 0.1 mg/mL, the HaCaT cell viability of the transdermal peptide recombinant type III collagen was higher than that of the commercially available or non-transdermal peptide recombinant type III collagen in all cases. The promotion effect of transdermal peptide recombinant type III collagen on the proliferation of HaCaT cells was better than that of general recombinant type III collagen (Figure 5).

**Figure 5.**
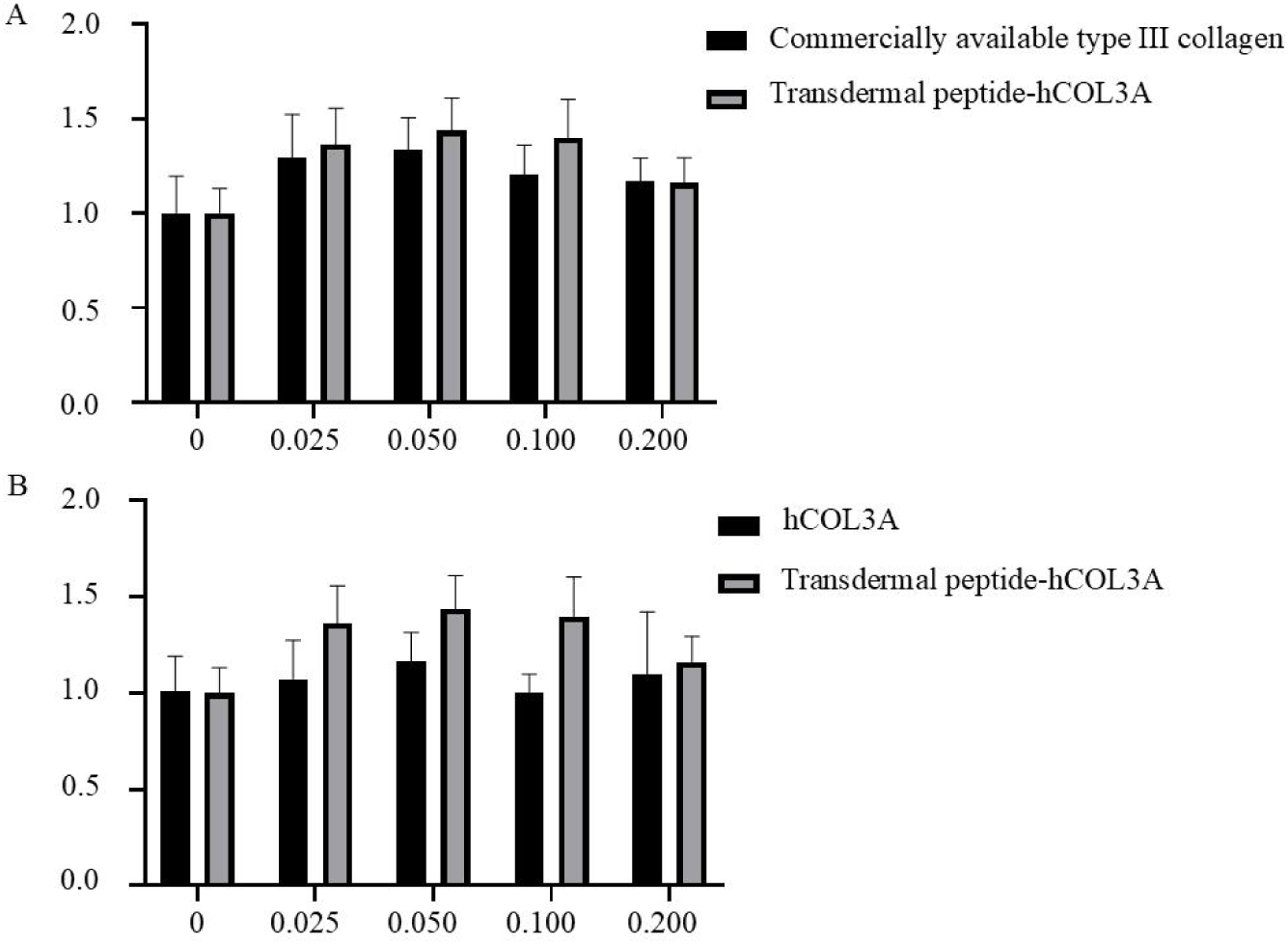
Cell proliferation experiment using HaCaT cells treated with recombinant type III collagen from different sources. (A) HaCaT cell proliferation experiment under the comparison of commercially available type III collagen with transdermal peptide recombinant type III collagen. (B) HaCaT cell proliferation assay without transdermal peptide recombinant type III collagen compared with transdermal peptide recombinant type III collagen

### 2.3 Transdermal peptide recombinant type III collagen hydrogel configuration

After examining the safety and biological activity of transdermal peptide recombinant type III collagen, we used it as the raw material to prepare the transdermal peptide recombinant type III collagen hydrogel, which included carbomer (thickening), propylene glycol (moisturizing), triethanolamine (pH regulation) and trehalose (stable protein structure), in order to adapt to the pH of human skin, we adjusted the ratio of each component (Table 2), and finally confirmed that 0.25% triethanolamine, 0.4% carbomer, 1% propylene glycol and 5% were used Trehalose could make the final pH of transdermal peptide recombinant type III collagen hydrogel 7.28 more suitable. Subsequently, the stability and hemolysis rate of transdermal peptide recombinant type III collagen hydrogel were detected.

**Table 2:**
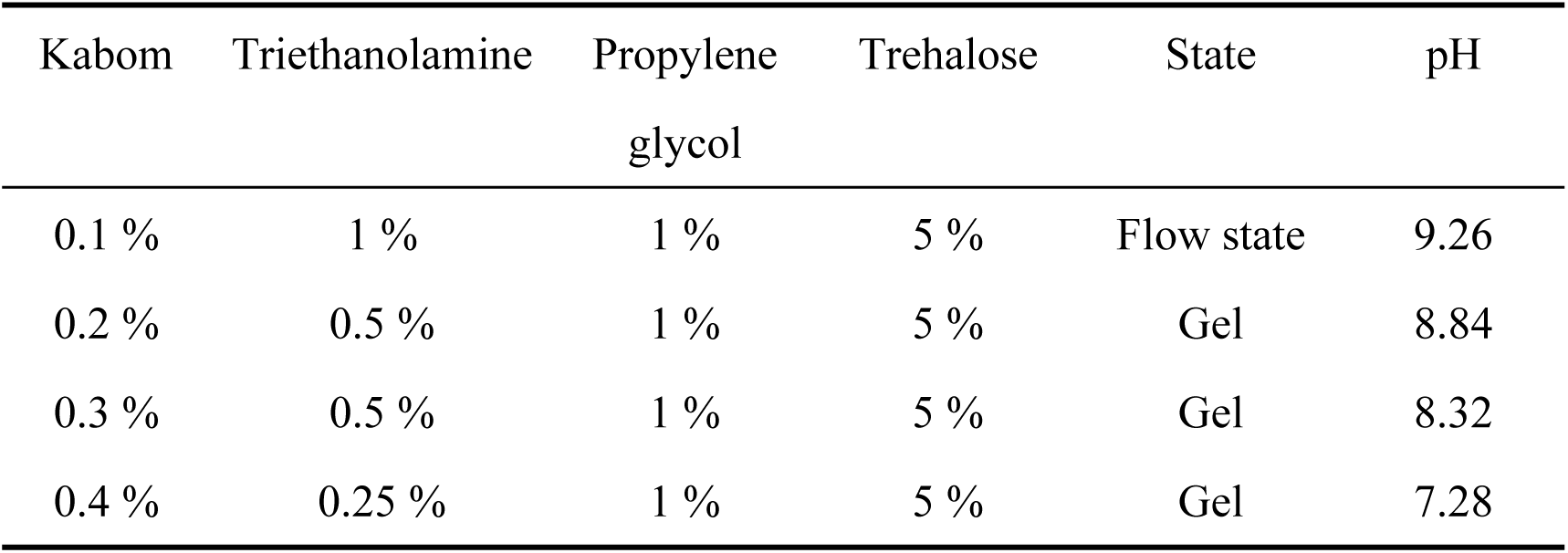
Optimization table of allocation ratio for each group (calculated by wt%)

| Kabom | Triethanolamine | Propylene glycol | Trehalose | State | pH |
| --- | --- | --- | --- | --- | --- |
| 0.1 % | 1 % | 1 % | 5 % | Flow state | 9.26 |
| 0.2 % | 0.5 % | 1 % | 5 % | Gel | 8.84 |
| 0.3 % | 0.5 % | 1 % | 5 % | Gel | 8.32 |
| 0.4 % | 0.25 % | 1 % | 5 % | Gel | 7.28 |

The prepared transdermal peptide recombinant type III collagen hydrogel was sealed and placed in an oven at 42±2 ℃, and sampled every day, and the SDS-PAGE results (Figure 6A) showed that the protein in the hydrogel was not degraded under the high temperature of 42±2 ℃, proving the stability of transdermal peptide recombinant type III collagen hydrogel. The hemolysis experimental data (Figure 6B) demonstrated the biosafety of transdermal peptide recombinant type III collagen hydrogel.

**Figure 6.**
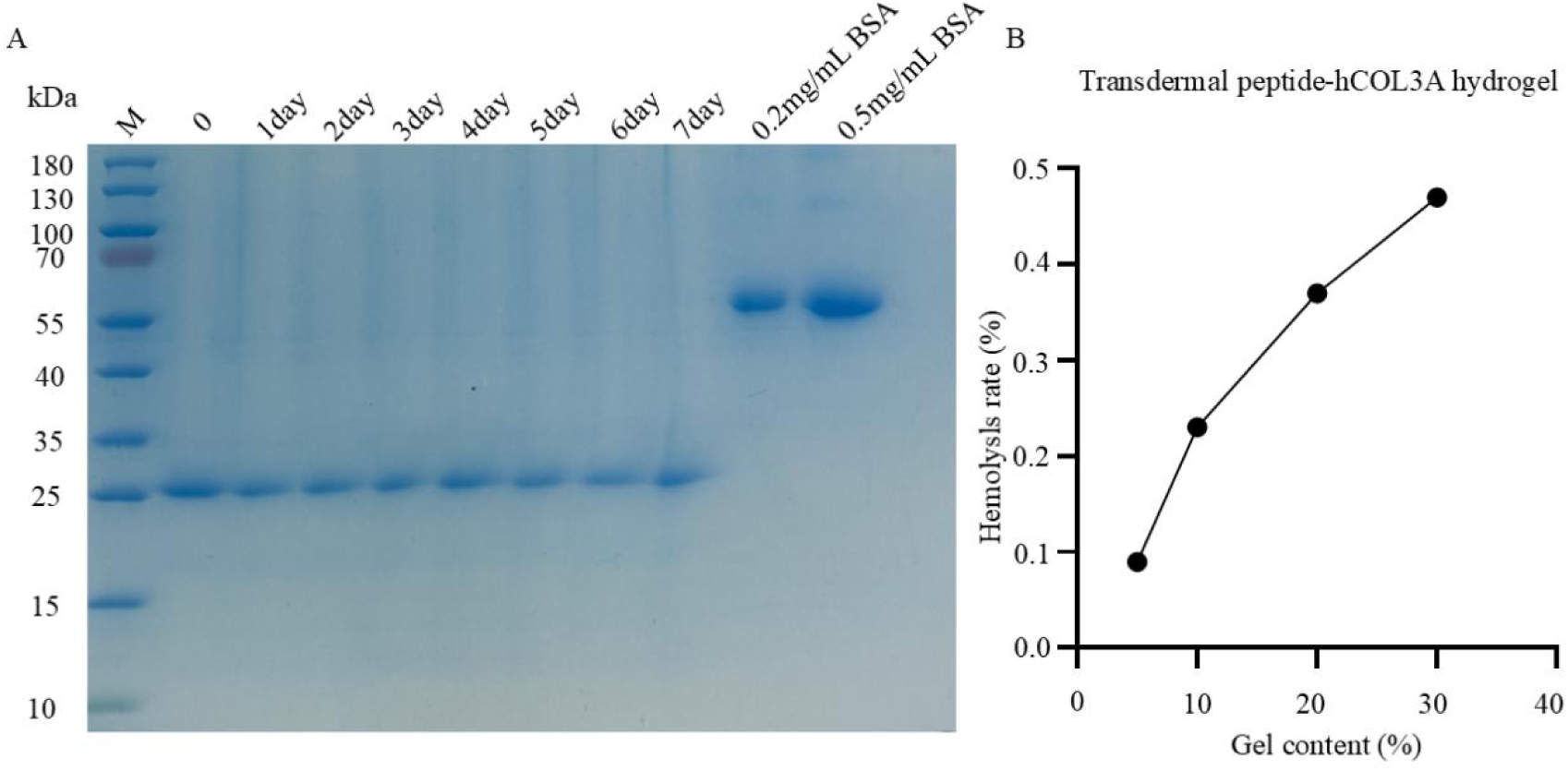
Stability of transdermal peptide recombinant type III collagen hydrogel at 42±2 °C). (A) SDS-PAGE analysis confirming no degradation of the collagen under high temperature conditions. (B) Hemolysis test demonstrating the biosafety of the hydrogel.

### 2.4 Biological activity of transdermal peptide recombinant type III collagen hydrogel

In order to prove that transdermal peptide recombinant type III collagen in the form of hydrogel does not lose the original proliferative effect of the protein, and the hydrogel itself had no toxic effect, we conducted multiple tests at both the cellular (HaCaT) and organismal (mouse) levels. In the HaCaT cell proliferation experiment controlled by transdermal peptide recombinant collagen type III hydrogel and hydrogel matrix (Figure 7A), we found that the hydrogel matrix did not cause toxic side effects to cells, and the results of the HaCaT cell proliferation experiment (Figure 7B) further proved that transdermal peptide recombinant type III collagen in hydrogel form also exhibited good biological activity.

**Figure 7.**
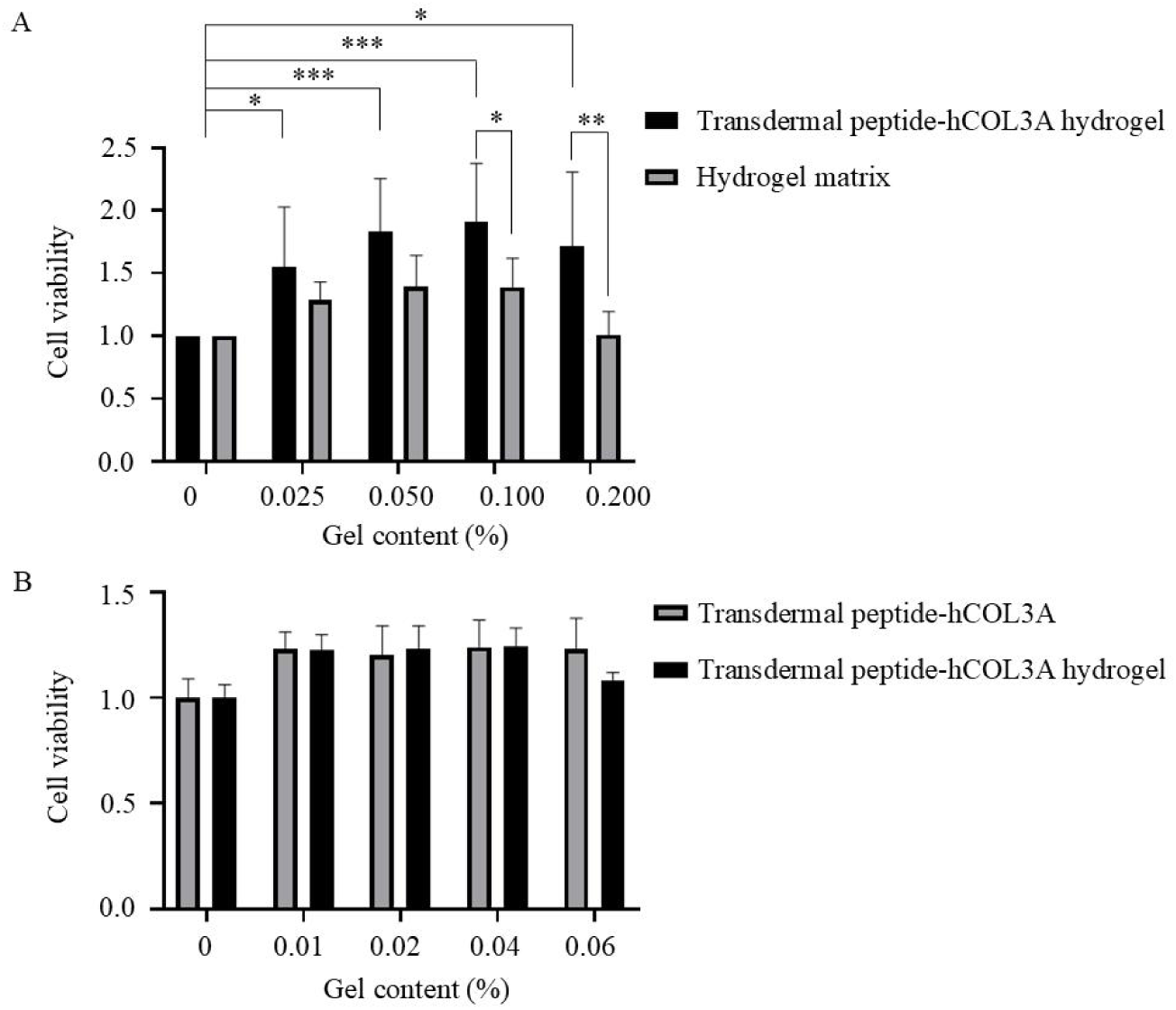
Proliferative effects of transdermal peptide recombinant type III collagen hydrogel on HaCaT cells. (A) Comparison of hydrogel matrix and collagen-loaded hydrogel (The data underwent T-test significance analysis, and its specific star-level correspondence is ⁎: 0.01<P<0.05, ⁎⁎: P<0.01, ⁎⁎⁎: P<0.001). (B) Cell proliferation results showing the hydrogel’s ability to promote HaCaT cell growth without toxic side effects.

At the individual level, we used a mouse skin wound healing model (Figure 9) to evaluate the wound healing time and status, and here we conducted experiments on female mice (Figure 8A) and male mice (Figure 8B), and the results showed that compared with the hydrogel matrix, transdermal peptide recombinant collagen type III hydrogel can promote wound healing faster, and has a wide range of gender adaptability, which can effectively promote trauma recovery in mice of different genders. Further, we tried to increase the protein concentration of transdermal peptide recombinant type III collagen in transdermal peptide recombinant type III collagen hydrogel, and observed mouse wound healing experiments. We found that when the protein concentration of transdermal peptide recombinant type III collagen was increased to 1 mg/mL, the healing effect of the gel on wound healing in mice was not significantly improved (Figure 9), indicating that the hydrogel has the ability to promote skin wound repair when a lower effective concentration of 0.2 mg/mL was used during the proportioning process.

**Figure 8.**
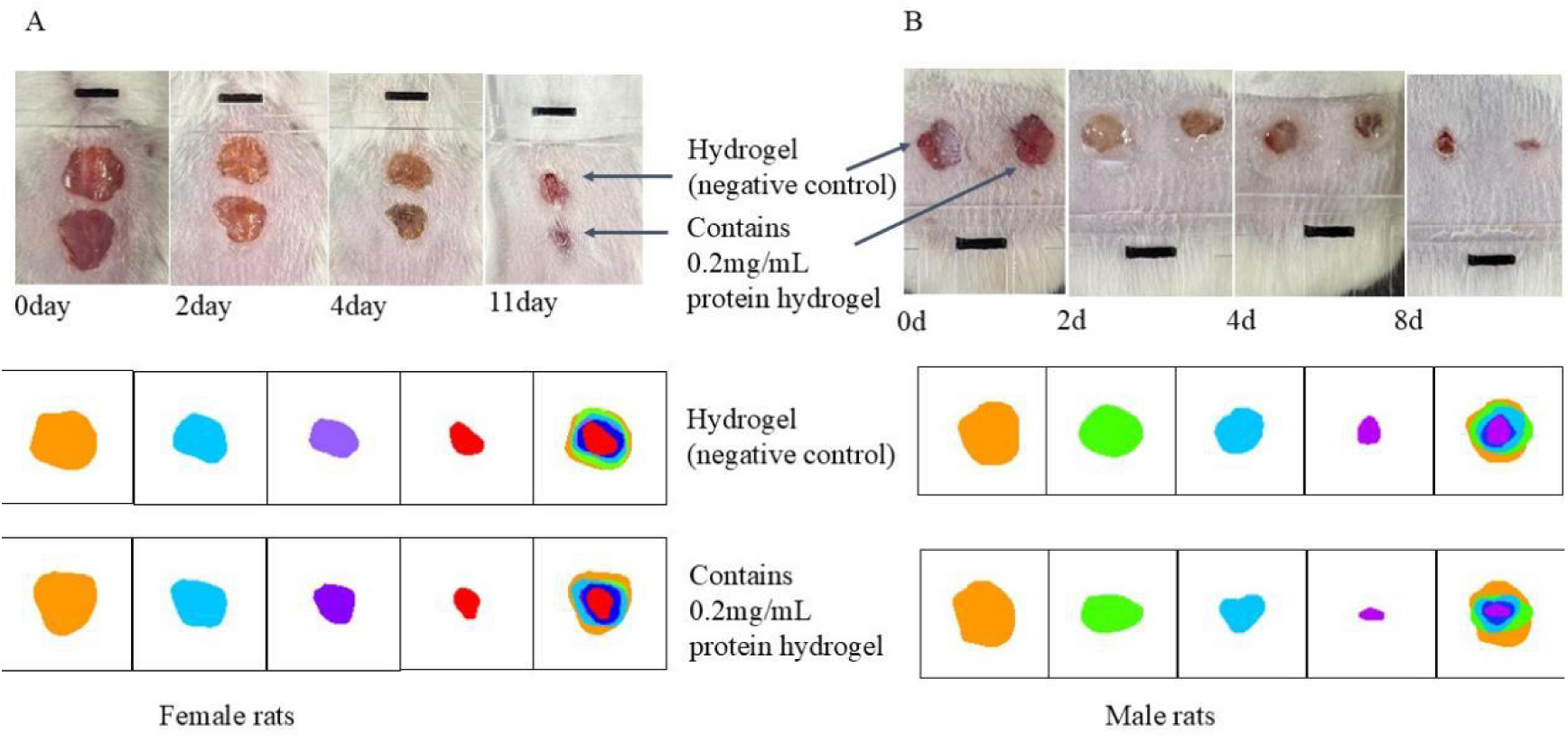
Mouse wound healing experiment using transdermal peptide recombinant type III collagen hydrogel. (A) Female mice wound healing status at 0.2 mg/mL collagen concentration. (B) Male mice wound healing status, showing faster recovery compared to the hydrogel matrix alone.

**Figure 9.**
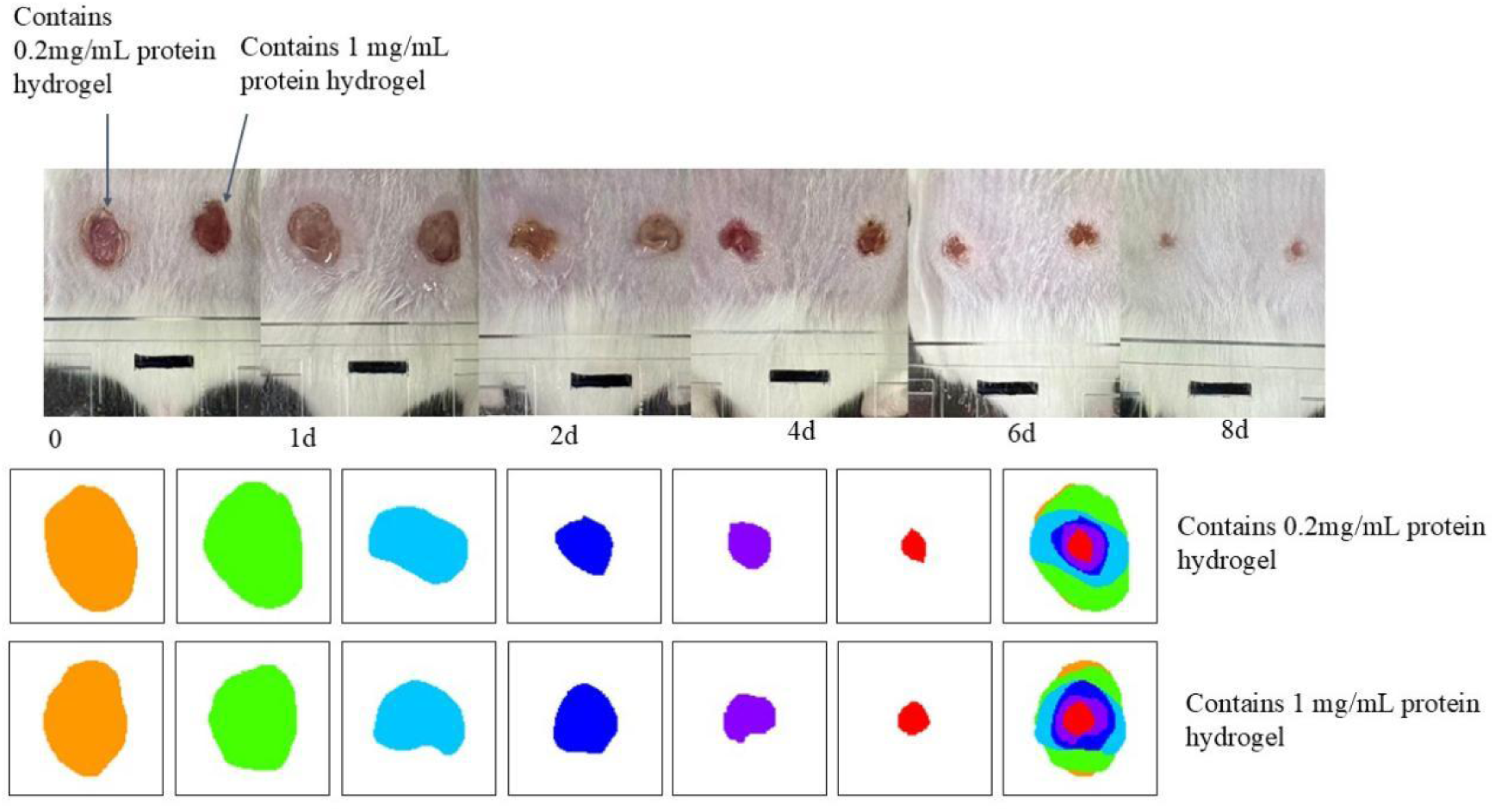
Comparison of wound healing effects of transdermal peptide recombinant type III collagen hydrogel at different concentrations. The healing effect at 0.2 mg/mL was optimal, showing significant improvement in wound closure compared to higher concentrations (1 mg/mL) and the protein-free hydrogel matrix.

### 2.5 Discussion

The results collectively demonstrate that transdermal peptide–recombinant type III collagen exhibits high purity, structural stability, and biological activity superior to conventional recombinant type III collagen. The hydrogel formulation not only preserves protein bioactivity but also provides enhanced thermal stability and hemocompatibility, enabling prolonged residence at wound sites.

Compared with traditional animal-derived collagen dressings, this engineered hydrogel offers several advantages:

Improved biocompatibility due to the absence of animal-derived pathogens and allergens.

Enhanced stability at physiological and elevated temperatures.

Superior wound healing performance at both cellular and organismal levels.

Beyond its advantages over traditional collagen dressings, the transdermal peptide-recombinant type III collagen hydrogel also exhibits marked improvements compared with other hydrogel systems reported in recent studies. Conventional collagen-based hydrogels—such as those crosslinked via disulfide bonds or modified with polyethylene glycol (PEG) and gelatin—often face challenges including rapid degradation, loss of mechanical integrity, and insufficient bioactivity under physiological stress. In contrast, the transdermal peptide incorporated in this study enhances both cutaneous penetration and protein stability, while maintaining a favorable microenvironment for cell proliferation and migration. The lamellar, porous microstructure observed under SEM further facilitates oxygen exchange and nutrient diffusion, critical for tissue regeneration.

When compared with synthetic polymer-based hydrogels (e.g., polyacrylamide or PVA-based systems), the present recombinant collagen hydrogel avoids potential cytotoxicity and inflammatory responses, while offering a biomimetic extracellular matrix–like architecture. Additionally, thermal and hemolysis assays confirm that the hydrogel maintains molecular integrity at elevated temperatures (42±2 ℃) and displays excellent blood compatibility, properties that surpass many existing hydrogel dressings that suffer from denaturation or reduced functionality under similar conditions.

At a functional level, the hydrogel demonstrated optimal wound healing efficacy at low collagen concentrations (0.2 mg/mL)-a significant advancement over typical collagen hydrogel formulations that often require higher loading to achieve comparable outcomes. This not only reduces production costs but also minimizes protein aggregation and potential immunogenic responses. The observed gender-independent wound healing efficacy further indicates the hydrogel’s robustness and universal applicability.

However, despite its encouraging results, certain limitations remain. The mechanical strength of the hydrogel under dynamic skin movement and its long-term degradation behavior in vivo warrant further evaluation. Moreover, the precise molecular mechanism underlying the synergistic effects of the transdermal peptide and recombinant collagen-such as their influence on cell signaling pathways related to angiogenesis and collagen remodeling—has yet to be fully elucidated.

## 3. Experimental steps

### 3.1 Molecular weight and purity analysis of transdermal peptide recombinant type III collagen

In the full-length sequence of human type III collagen, an active fragment that had not been reported and studied so far was intercepted, and General Biology (Anhui) Co., Ltd. was entrusted to synthesize the gene and construct it on the pPIC9K vector, and the amino acid sequence of transdermal peptide recombinant type III collagen was (SEQ ID NO.1):

FEGPASRPGFPGMKGHRGFDGRNGEKGETGAPGLKGENGLPGENGAPGPMGPRGAPGERGRPGLPGAAGARGNDGARGSDGQPGPPGPPGTAGFPGSPGAKGEVGPAGSPGSNGAPGQRGEPGPQGHAGAQGPPGPPGINGSPGGKGEMGPAGIPGAPGLMGARGPPGPAGANGAPGLRGGAGEPGKNGAKGEPGPRGERG EAGIPGV PGAKGEDGKDGSPGEPGANG

Transdermal peptide amino acid sequence (SEQ ID NO.2):

FEGPASR

Collagen amino acid sequence (SEQ ID NO.3):

PGFPGMKGHRGFDGRNGEKGETGAPGLKGENGLPGENGAPGPMGPRGAPGERGRPGLPGAAGARGNDGARGSDGQPGPPGPPGTAGFPGSPGAKGEVGPAGSPGSNGAPGQRGEPGPQGHAGAQGPPGPPGINGSPGGKGEMGPAGIPGAPGLMGARGPPGPAGANGAPGLRGGAGEPGKNGAKGEPGPRGERGEAGIPGVPG AKGED GKDGSPGEPGANG

The recombinant plasmid containing SEQ ID NO.1 sequence was transformed into the *Pichia pastoris* GS115 strain by the chemical transformation method. The expressible engineered bacteria were screened, and the expressible engineered strain was used to express transdermal peptide recombinant type III collagen, and the fermentation broth was collected for subsequent purification.

The transdermal peptide recombinant type III collagen purified by Q and SP ion-exchange chromatography was reconstituted, dialyzed to 25 mmol/L NaCl, 10 mmol/L Tris pH 7.2, and the samples were concentrated to 2-2.5 mg/mL through a 10 kDa ultrafiltration concentrate tube for freeze-drying. The freeze-dried sample was reconstituted in PBS buffer and centrifuged at 13,000 rpm for 1 h at 4 ℃. After dilution to an appropriate concentration, the sample was mixed with MBP and RNase proteins, followed by another centrifugation step under the same conditions (13,000 rpm, 1 h, 4 ℃). The mixture was subsequently analyzed using a Superdex™ 200 Increase 10/300 GL gel filtration chromatography column. The freeze-dried transdermal peptide recombinant type III collagen was sent to the scientific research dog instrument test platform for testing, and a 1 μL sample was dried naturally, and then 1 μL matrix was added to air dry. The instrument was performed using matrix-assisted laser desorption/ionization time-of-flight mass spectrometry (MALDI-TOF; Ultraflextreme model). The matrix used was α-cyano-4-hydroxycinnamic acid (CHCA), the test mode was positive spectrum, the laser energy was 100%, and the test molecular weight range was 10-100 kDa. In addition, transdermal peptide recombinant type III collagen purified by an ion exchange column and freeze-dried transdermal peptide type III. was dissolved using dd H2O, frozen, centrifuged at 13,000 rpm at 4 ℃, and analyzed by high-performance liquid chromatography using Shimadzu LC-2030 Plus. Mobile phase A: ultrapure water: trifluoroacetic acid (TFA) = 99.9 %: 0.1 %, mobile phase B: methanol: ultrapure water: TFA = 80 %: 19.9 %: 0.1 %, transdermal peptide recombinant type III collagen was loaded with 10 μL for gradient elution; Analysis was performed using Shimadzu LC-2030 Plus data processing software.

### 3.2 Circular Dichroism (CD) Spectroscopic Analysis of Transdermal Peptide–Recombinant Type III Collagen

Lyophilized transdermal peptide–recombinant type III collagen, purified by ion-exchange chromatography, was dissolved in two buffer systems: Buffer A (135 mmol/L NaCl, 2.7 mmol/L KCl, 10 mmol/L PBS, pH 7.2) and Buffer B (135 mmol/L NaCl, 2.7 mmol/L KCl, 10 mmol/L PBS, pH 6.2). Following overnight dialysis at 4 ℃ the supernatant was concentrated using a 10 kDa ultrafiltration tube, and protein samples were prepared at a final concentration of 10 µM. CD spectra were recorded on a Chirascan qCD spectrometer over the wavelength range of 190–240 nm, with each sample scanned in triplicate. Buffer baselines were subtracted during data processing.

### 3.3 Microscopic morphology analysis of the surface of transdermal peptide recombinant type III collagen raw materials

The silicon substrate was cleaned with anhydrous ethanol, coated with conductive glue, the lyophilized transdermal peptide recombinant type III collagen was spread evenly on the conductive layer, and gently blown off the non-adherent proteins with an air stream. The sample was gold-plated and imaged using a Hitachi cold field emission scanning electron microscope to observe its surface micromorphology. Scanning electron microscopy was further used to examine the structure of the transdermal peptide recombinant type III collagen at magnifications of 200× and 400×.

### 3.4 Transdermal peptide recombinant type III collagen in vitro metamorphic skin reaction

The guinea pigs used in the test were healthy, newly adult guinea pigs, and their weight should be 300 g∼500 g at the beginning of the test, regardless of male or female, such as female rats should be nulliparous or non-pregnant. Before the experiment, the hair on the left side of the guinea pig’s back was shaved one day in advance, with a bare skin range of about 4 cm × 4 cm for the first provocation administration after 24 h.

The lyophilized transdermal peptide recombinant type III collagen was dissolved in PBS and filtered and sterilized using a 0.22 μm filter head. Administration was carried out on the shaved left side of the guinea pig’s back: 1 mL of transdermal peptide recombinant type III collagen was applied at a concentration of 0.2 mg/mL, and after the protein solution was dried, the administration site was fixed with sterile gauze for 6 h, during which the state of the guinea pig was observed. The skin condition of the administration site was recorded after 24±2 h and 48±2 h after administration, respectively.

### 3.5 Permeation experiment of recombinant type III collagen on excised mouse skin between female and male

#### Skin preparation

The abdominal skin of ICR mice was excised, and hair, fat, and connective tissue were removed. The skin was rinsed with distilled water and saline, then stored at 4 ℃ for later use.

#### Device assembly

The rat skin was fixed between the donor and receptor chambers of the diffusion cell. The receptor chamber was filled with 15 mL of PBS preheated to 32 ℃ and maintained at a constant temperature of 32 ℃ with stirring at 100 r/min.

#### Administration

A 1 mL aliquot of the collagen sample was added to the donor chamber (experimental group: Transdermal peptide recombinant Type III Collagen; control group: non-transdermal peptide recombinant type III collagen).

#### Sampling and detection

Samples (0.2 mL) were collected from the receptor chamber at 1, 2, 4, 6, and 8 hours and immediately replenished with an equal volume of isothermal PBS. Protein concentrations were determined for each sample.

#### Data analysis

The cumulative permeation per unit area was calculated using the following equation.

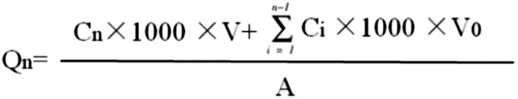

where Cn-collagen concentration measured at the nth sampling point; Collagen concentration measured at Ci-ith sampling point; 1000 - unit conversion factor; V-Acceptance cell volume, which was 15 mL; V0 - sample volume, which was 0.2 mL; A-permeation area, which was 1.13 cm^2^.

### 3.6 Cell proliferation experiment

#### HaCaT cell preparation

HaCaT cells were resuscitated and passaged using complete medium containing 10% fetal bovine serum and 1% penicillin–streptomycin bispecific antibody. The cells were cultured for approximately 3 days per generation. After three stable passages, the cells were counted, diluted to a concentration of 1×10⁵ cells/mL, and seeded into 96-well plates at 100 μL per well. The plates were incubated at 37 °C for 24 h in a cell culture incubator for subsequent experiments.

#### Experimental groups

Group 1: Commercially available recombinant type III collagen and transdermal peptide recombinant type III collagen

Group 2: No transdermal peptide recombinant type III collagen and transdermal peptide recombinant type III collagen

#### Treatment concentrations

The protein concentrations were set to 0, 0.025 mg/mL, 0.05 mg/mL, 0.1 mg/mL and 0.2 mg/mL.

Group 3: Transdermal peptide recombinant type III collagen hydrogel and transdermal peptide recombinant type III collagen raw materials.

Group 4: Transdermal peptide recombinant type III collagen hydrogel and hydrogel matrix.

#### Treatment concentrations

The protein concentrations were set to 0, 0.025 mg/mL, 0.05 mg/mL, 0.1 mg/mL, and 0.2 mg/mL.

For the hydrogel groups, concentrations of 5%, 10%, 20%, 30%, and 50% were prepared, corresponding to effective protein concentrations of 0.01, 0.02, 0.04, and 0.06 mg/mL. The same concentration of pure transdermal peptide recombinant type III collagen was used as the control group.

For the hydrogel-matrix group, the hydrogel matrix without protein served as the blank carrier control, and the culture medium served as the blank control. Each group was prepared in six replicate wells, and all solutions were sterilized using a 0.22 μm filter membrane.

#### Experimental procedure

During the experiment, the medium in each 96-well plate was aspirated, and 100 μL of the diluted experimental or control sample was added to each well. The plates were then incubated again at 37 °C for 24 h prior to CCK-8 detection.

#### CCK-8 assay configuration

The CCK-8 reagent was prepared by adding 5 μL of CCK-8 solution to every 95 μL of medium and mixing thoroughly. After incubation, the PBS and medium in the wells were aspirated, and 100 μL of the reaction mixture was added to each well. The absorbance at 450 nm was measured using a microplate reader after 30 min of incubation at 37 °C. Cell viability was calculated based on the obtained absorbance values.3.7 In vitro trauma recovery experiment of transdermal peptide recombinant type III collagen hydrogel

#### Animals and anesthesia

6∼8 week-old Kunming mice were selected and randomly assigned into groups. Anesthesia was administered via ether inhalation, and the loss of the righting reflex was used as the end point of anesthesia.

#### Trauma model establishment

After confirming that anesthesia was sufficient, the dorsal hair of the mice was shaved and the skin was disinfected with 75% ethanol. The limbs were immobilized, and two full-thickness wounds of similar size were created on the exposed dorsal skin using a sterilized 5 mm biopsy punch for experimental treatment compared with the control.

#### Experimental grouping and administration

Experimental Group 1: Hydrogel containing transdermal peptide recombinant type III collagen (effective concentration 0.2 mg/mL)

Experimental group 2: Transdermal peptide recombinant type III collagen hydrogel (effective concentration 1 mg/mL)

Control group: Protein-free hydrogel matrix (blank carrier)

Administration method: An equal amount of hydrogel (0.05 g) was applied evenly to the wound surface.

#### Trauma recovery monitoring

The mice were fixed in the supine position, and wound healing was recorded at each time point. Photographs were taken to document the morphological changes and the healing process of the wounds. Wound healing was analyzed using ImageJ image analysis software.

## Author Contributions

Prof. Jingjun Hong collected the financial funding, top designed and supervised the whole procedure of research and paper writing. Yue Liu was responsible for the allergenicity testing of the raw material, hydrogel preparation, cell proliferation assays, stability testing, and the mouse wound healing study, and drafted the corresponding sections of the manuscript. Wansen Tan and Prof. Jingjun Hong performed the analysis of raw material purity and molecular weight, conducted circular dichroism (CD) spectroscopy, scanning electron microscopy (SEM), and the mouse skin permeation assay, and drafted the respective sections. The preparation of the transdermal peptide recombinant type III collagen and non-transdermal peptide recombinant type III collagen material were done by Wansen Tan and Yue Liu, respectively.

## Acknowledgments

This research was financially supported by the Natural Science Foundation of Hefei City (HZR2422), the Research Foundation of Anhui Zhongsheng Anlan Health Industry Co., Ltd (K160162428), the National Natural Science Foundation of China (grant 31970669), and the Scientific Research Foundation for Returned Scholars, Anhui University (S020318006/067). We thank Yunru Yang and Dr. Tengchuan Jin (USTC) for their help with CD data collection. We thank the core facility of School of Life Science and Medical Engineering, Anhui University.

## Ethical Statement

All animal experiments conducted in this study were performed in strict accordance with the ethical guidelines and regulations set forth by the Institutional Animal Care and Use Committee (IACUC) of Anhui University. Approval Number: IACUC (AHU)-2025-005.

